# The N-Glycome regulates the endothelial-to-hematopoietic transition

**DOI:** 10.1101/602912

**Authors:** Dionna M. Kasper, Jared Hintzen, Yinyu Wu, Joey J. Ghersi, Hanna K. Mandl, Kevin E. Salinas, William Armero, Zhiheng He, Ying Sheng, Yixuan Xie, Daniel W. Heindel, Eon Joo Park, William C. Sessa, Lara K. Mahal, Carlito Lebrilla, Karen K. Hirschi, Stefania Nicoli

**Affiliations:** Yale Cardiovascular Research Center, Department of Internal Medicine, Section of Cardiology, Yale University School of Medicine, New Haven, CT 06511, USA; Department of Genetics, Yale University School of Medicine, New Haven, CT 06510, USA; Department of Pharmacology, Yale University School of Medicine, New Haven, CT 06510, USA; Vascular Biology & Therapeutics Program, Yale University School of Medicine, New Haven, CT 06520, USA; Biomedical Chemistry Institute, Department of Chemistry, New York University, New York, NY 10003, USA; Department of Chemistry, University of Alberta, Edmonton, AB, T6G 2G2, Canada; Department of Chemistry, University of California, Davis, CA 95616; Developmental Genomics Center, Cell Biology Department, University of Virginia School of Medicine, Charlottesville, VA 22908

## Abstract

Hematopoietic stem and progenitor cells (HSPCs) that establish and maintain the blood system in adult vertebrates arise from the transdifferentiation of hemogenic endothelial cells (hemECs) during embryogenesis. This endothelial-to-hematopoietic transition (EHT) is tightly regulated, but the mechanisms are poorly understood. Here, we show that microRNA (miR)-223-mediated regulation of N-glycan biosynthesis in endothelial cells (ECs) regulates EHT. Single cell RNA-sequencing revealed that miR-223 is enriched in hemECs and in oligopotent nascent HSPCs. miR-223 restricts the EHT of lymphoid/myeloid lineages by suppressing the expression of mannosyltransferase *alg2* and sialyltransferase *st3gal2*, two enzymes involved in N-linked protein glycosylation. High-throughput glycomics of ECs lacking miR-223 showed a decrease of high mannose versus sialylated complex/hybrid sugars on N-glycoproteins involved in EHT such as the metalloprotease Adam10. Endothelial-specific expression of an N-glycan Adam10 mutant or of the N-glycoenzymes phenocopied the aberrant HSPC production of miR-223 mutants. Thus, the N-glycome plays a previously unappreciated role as an intrinsic regulator of EHT, with specific mannose and sialic acid modifications serving as key endothelial determinants of their hematopoietic fate.

**One Sentence Summary:** The N-glycan “sugar code” governs the hematopoietic fate of endothelial cells and regulates blood stem cell production in vivo.

## Main Text

In vertebrates, HSPCs with long-term engraftment capability are specified during embryogenesis by the transdifferentiation of ECs within the aorta-gonad-mesonephros (AGM) region of the dorsal aorta (*1*–*3*). This EHT occurs in hemECs, a subset of ECs that co-express vascular and hematopoietic genes. The precise interplay between multiple signaling cascades (*1, 2, 4*) enables progressive loss of EC gene expression and concomitant increase in hematopoietic potential, with hemECs giving birth to nascent HSPCs (*2, 3*). HSPCs then delaminate from the vascular wall into the circulation to colonize secondary hematopoietic organs, where they proliferate and differentiate to generate all blood cells throughout life (*5*–*7*). The EC and/or hemEC determinants that regulate EHT and thus the generation of HSPCs is incompletely understood.

Here, we investigated the function of miR-223, for which genetic deletion (miR-223Δ/Δ) results in excess nascent HSPCs in the zebrafish AGM (*8*). However, the mechanisms underlying miR-223 function in EHT are unknown (*8*–*11*).

To examine miR-223 expression during EHT, we generated a zebrafish transgenic reporter. We integrated the Gal4 transcriptional activator (*12*) downstream of the endogenous miR-223 promoter (miR-223:Gal4) without interfering with miRNA expression, and coupled it with *Tg(5XUAS:eGFP)*^*nkuasgfp1a*^ and *Tg(kdrl:hras-mCherry)*^*s896*^ reporters so that miR-223-expressing ECs are labeled with both GFP and mCherry (miR-223:GFP+ kdrl:mCH+) (Fig. 1A and fig. S1A). Endogenous miR-223 was enriched in miR-223:GFP+ kdrl:mCH+ cells versus miR-223:GFP-kdrl:mCH-cells (fig. S1B), as expected. Moreover, re-expression of miR-223 from its endogenous promoter in miR-223:Gal4+ cells rescued the overexpansion of *cmyb+* HSPCs in miR-223Δ/Δ(fig. S1C). Thus, miR-223:GFP+ cells report the endogenous expression and function of miR-223.

**Fig 1.**
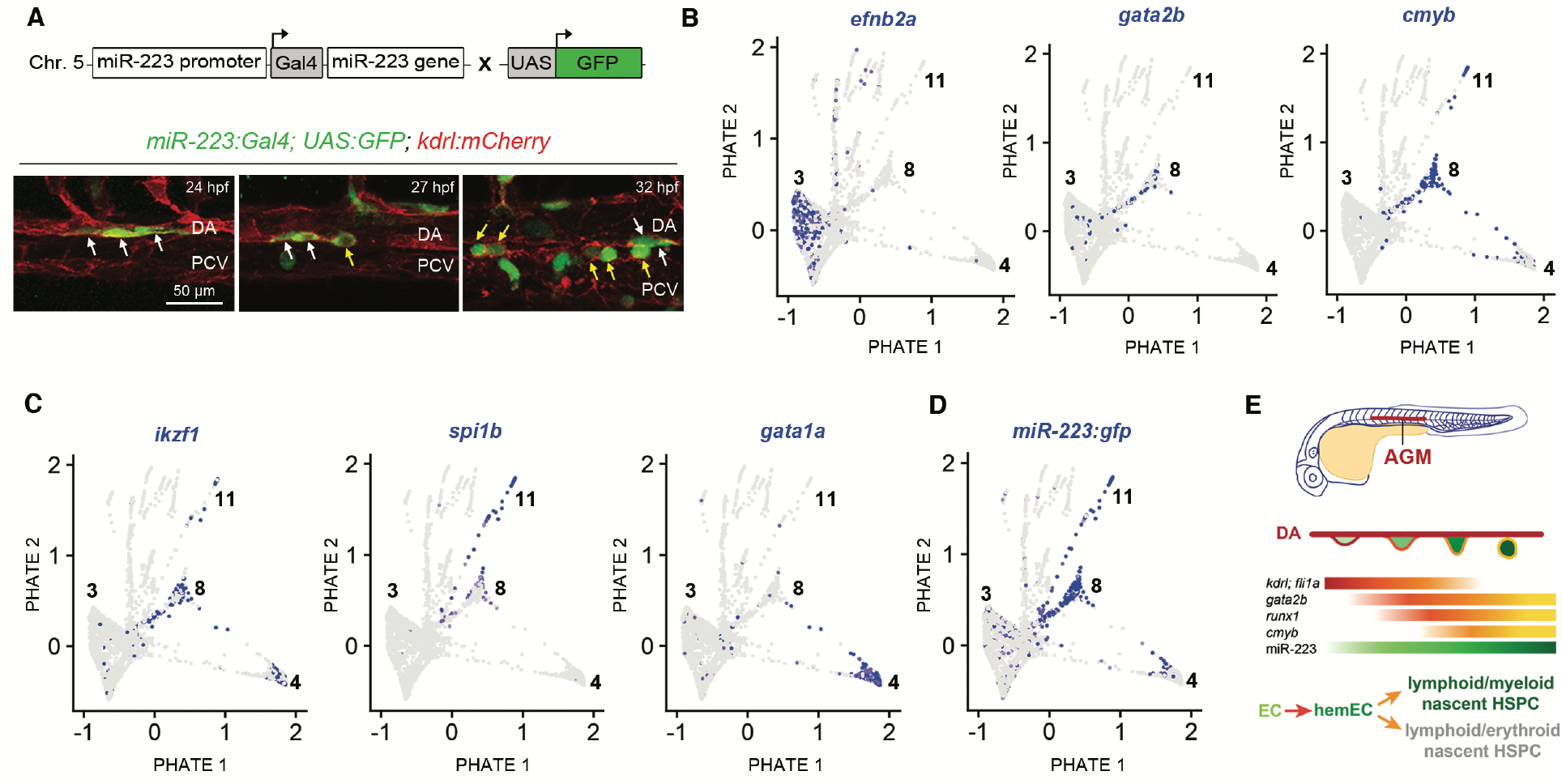
miR-223 is expressed in hemECs undergoing EHT. **(A)** Lateral Z-projections of the zebrafish AGM. White and yellow arrows point to flat and budding miR-223:GFP+ kdrl:mCH+ cells in the VDA, respectively. **(B-D)** scRNA-seq PHATE plots of 27 hpf wild-type kdrl:mCH+ trunk ECs showing relative expression levels of EHT (**B**) blood lineage (**C**), and *miR-223:gfp* (**D**) genes (n = 6227 wild-type cells). Arterial (branch 3) and EHT (branch 8, 11, 4) PHATE trajectories are indicated. **(E)** Overview of zebrafish EHT. In the AGM, a flattened EC (red outline) gains hemogenic potential (orange outline), and buds as a nascent oligopotent HSPC (yellow outline) from the DA wall. Red to yellow gradients depict the expression patterns of endothelial and EHT markers and are colored based on their expression in indicated cell types. The green gradient and text represent the extent of miR-223 expression during EHT. Abbreviations: DA, dorsal aorta; PCV, posterior cardinal vein

miR-223:GFP+ kdrl:mCH+ cells were dispersed in a salt and pepper pattern, mostly within the AGM (Fig. 1A and fig. S1, D and E). The AGM population of miR-223:GFP+ kdrl:mCH+ cells increased from the onset to the peak of EHT, at 24 to 32 hours post fertilization (hpf) (fig. S1E). In addition, they manifested a heterogeneous pattern of flat to bulging morphologies, and underwent delamination from the AGM (Fig. 1A, fig. S1, F and G, and Movie S1). miR-223:GFP+ kdrl:mCH+ cells showed elevated expression of endothelial (*kdrl*) and EHT markers, including *gata2b* and *runx1* in hemECs (*13*) and *cmyb* in nascent HSPCs (*5*) (fig. S1H). Accordingly, nascent HSPCs showed elevated expression of endogenous mature miR-223 (fig. S1I), suggesting that miR-223 is expressed in ECs undergoing EHT.

To further discern the distinct molecular subtypes among miR-223-expressing ECs within the AGM, we performed single cell RNA-sequencing (scRNA-seq). Briefly, we isolated kdrl:mCH+ ECs from the trunks of miR-223:GFP+ kdrl:mCH+ zebrafish embryos during early EHT (fig. S2A). To reveal gradual transitions through EC differentiation, we analyzed scRNA-seq data with PHATE (*14*). PHATE is a dimensionality reduction method that simultaneously denoises transcriptome data and organizes cells into a tree hierarchy based on the cell-to-cell gene expression continuums.

Kdrl:mCH+ ECs formed a vascular tree composed of 14 different branches. These branches corresponded to embryonic specification trajectories, identified by the expression of known markers (fig. S2, A and B and Data S1). Importantly, we observed one *efnb2a+* arterial trajectory (branch 3) from which cells start to co-express a continuum of early and late EHT markers, including *gata2b and runx1* in hemECs and *cmyb* in nascent HSPCs (branch 8, Fig. 1B, fig. S2B, and Data S1). This branch split into two trajectories that included *cmyb+* nascent HSPCs expressing early primed markers for the lymphoid/myeloid lineages (branch 11, marked by *ikzf1/spi1b*) or lymphoid/erythroid lineages (branch 4, marked by *ikzf1/gata1a*) (Fig. 1C, fig. S2B, and Data S1). Thus, our data suggest that oligopotent nascent HSPCs are produced during EHT in the AGM.

To confirm that these trajectories were associated with EHT, we performed gene ontology (GO) analysis of the branch defining genes. GO classifications showed that as cells progress through clusters 3, 8, 11 and 4, they lose vascular development terms while gaining hematopoietic, protein biosynthesis and N-glycosylation terms (*15*). Cell migration and cell cycle transcripts were mostly acquired at the branch points 11 and 4 where the primed nascent HSPCs likely begin to delaminate and/or proliferate (fig. S2C and Data S2) (*1, 2*).

Next, we analyzed kdrl:mCH+ cells expressing *miR-223:gfp* within the identified EHT trajectories. We found that ECs expressing *gfp* transcripts comprised ~73 to 100% of branches 8 (hemECs and nascent HSPCs) and 11 (lymphoid/myeloid-primed HSPCs), and only ~17 % of branch 4 (lymphoid/erythroid-primed HSPCs) (Fig. 1D and fig. S2, D and E). Consistent with endogenous miR-223 expression, scRNA-seq average gene expression revealed that *miR-223:gfp+* ECs were enriched in EHT markers, and can express both lymphoid/myeloid- and lymphoid/erythroid-primed HSPC lineage markers (fig. S2F and Data S1). Together, these analyses suggest that miR-223 is enriched in hemECs and nascent HSPCs during the EHT of oligopotent lymphoid/myeloid-primed HSPCs (Fig. 1E).

Next, we analyzed miR-223Δ/Δ zebrafish for EHT defects. Compared to wild-type, miR-223Δ/Δ embryos displayed an increase in gata2b:GFP+ or runx1:GFP+ hemECs in the AGM (Fig. 2, A and B, and fig. S3A). As observed previously, cmyb:GFP+ kdrl:mCH+ nascent HSPCs were increased at 32 and 36 hpf in miR-223Δ/Δ(Fig. 2C) (*8*). Together, these data indicate that miR-223Δ/Δembryos have an excess of both hemECs and nascent HSPCs.

**Fig. 2.**
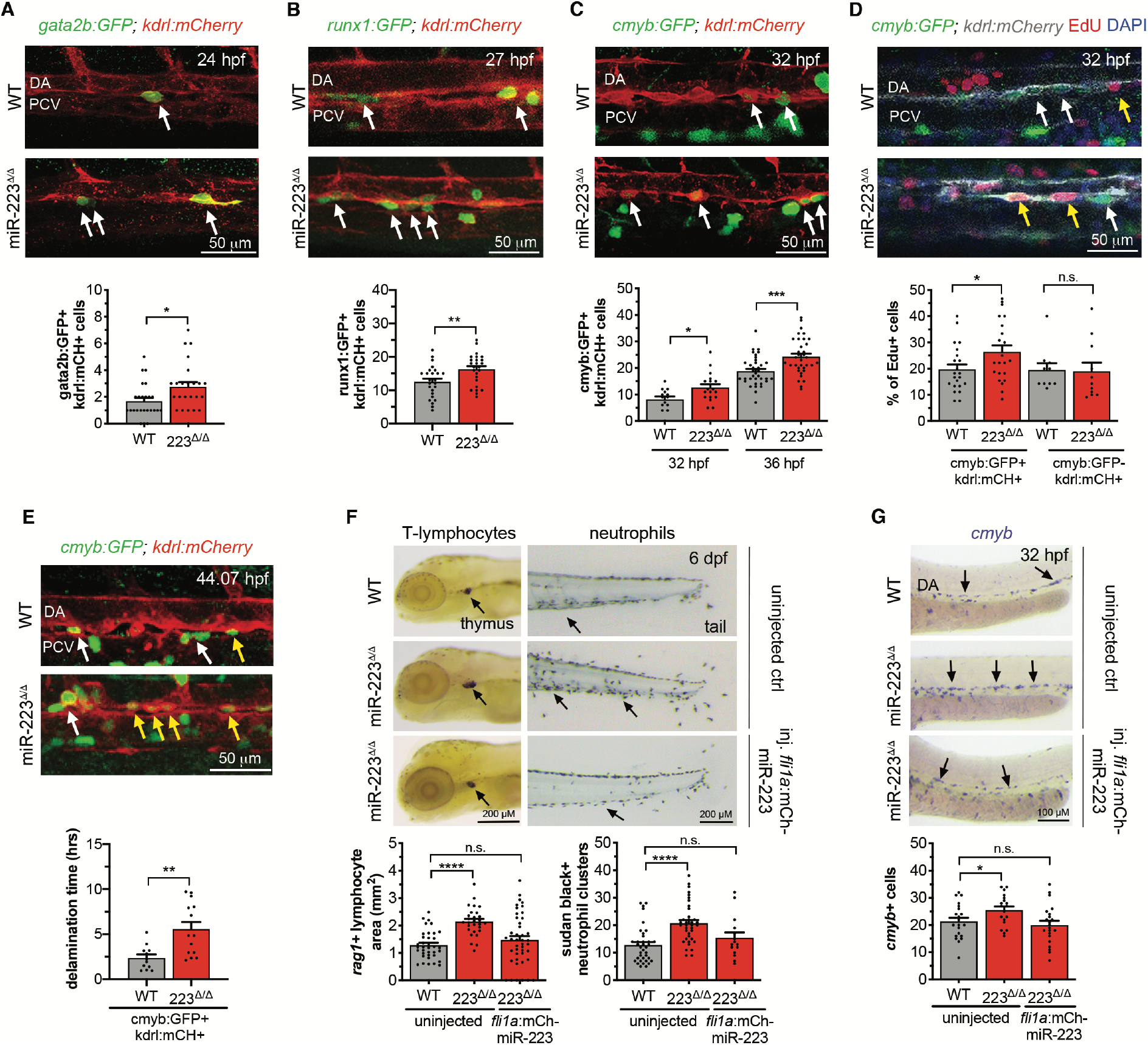
miR-223 is a conserved endothelial inhibitor of EHT and lymphoid/myeloid-oligopotent HSPC production. **(A-C)** *top*, Lateral Z-projections of the zebrafish AGM. White arrows point to gata2b:GFP+ (**A**) or runx1:GFP+ (**B**) kdrl:mCH+ hemECs or cmyb:GFP+ kdrl:mCH+ nascent HSPCs (**C**). *bottom*, Mean ± SEM number of gata2b:GFP+ kdrl:mCH+ cells (n = 24 - 25 embryos) (**A)**, runx1:GFP+ kdrl:mCH+ cells (n = 25 embryos) (**B)**, or cmyb:GFP+ kdrl:mCH+ cells (n = 13-18, 32 hpf; n= 34-36, 36 hpf) (**C**). 36 hpf data was previously published (*8*). (**D**) *top*, Yellow and white arrows point to EdU+ and EdU− nascent HSPCs, respectively. *bottom*, Mean ± SEM percentage of EdU+ nascent HSPCs (n = 22 embryos) or ECs (n = 10 – 11 embryos). (**E**) *top*, Yellow arrows in timelapse Z-projection point to nascent HSPCs embedded within the zebrafish AGM and white arrows indicate HSPCs that have bud into the subaortic space. *Bottom*, Mean ± SEM of delamination time for nascent (n = 17 cells). **(F-G)** *top*, Representative T-lymphocyte marker *rag1* expression, Sudan black neutrophil staining (**F**), and *cmyb* expression (**G**). *bottom*, Mean ± SEM area of *rag1* expression (n = 18 – 35 embryos), Sudan Black+ clusters (n = 36 - 37 embryos) (**F**), and *cmyb+* cell number (n = 18 – 21 embryos) (**G**). The *fli1a:mCH-miR-223* construct was injected to rescue miR-223 expression in miR-223Δ/Δ ECs as described in fig. S3I. Significance calculations are represented as not significant (n.s.) p > 0.05, ^*^ p ≤ 0.05, ^* *^ p ≤ 0.01, ^* * *^ p ≤ 0.01, ^* * * *^ p ≤ 0.0001, unpaired, two-tailed Mann-Whitney U test. Abbreviations as in Fig. 1, and KM, kidney marrow.

We examined whether miR-223 function is conserved in mammals by analyzing EHT in mice. miR-223 was abundantly expressed in ECs and hemECs compared to non-ECs isolated from mouse embryonic day (e) 10.5 AGM, the primary site of definitive HSPC production in mice (*2, 6*) Compared to wild-type mice, mice with global miR-223 knockout (KO) (*9*) displayed elevated hemECs in the e10.5 AGM and HSPCs at secondary hematopoietic sites (fig. S3, D and E). These results suggest that miR-223 is a conserved inhibitor of hemEC and HSPC production.

Next, we determined if nascent HSPCs in miR-223Δ/Δ zebrafish have altered behaviors. First, we examined their proliferative capacity by an EdU incorporation assay. We observed an increase in the fraction of EdU+ nascent HSPCs in miR-223Δ/Δ versus controls, whereas the fraction of EdU+ ECs was similar between genotypes (Fig. 2D).

Additionally, we used in vivo time-lapse imaging to quantify the delamination of nascent HSPCs from the AGM. Interestingly, delamination times were significantly longer (2.3 ± 0.9 hours) for miR-223Δ/Δcompared to wild-type nascent HSPCs (Fig. 2E, fig. S3F, and Movies S2 and S3). Although slow, *cmyb+* HSPCs eventually delaminated and were still expanded in secondary hematopoietic organs of miR-223Δ/Δ embryos, namely in the caudal hematopoietic tissue (CHT, the mammalian fetal liver equivalent) at 2.5 days post fertilization (dpf) (*8*), as well as, in the thymus and kidney marrow (the mammalian bone marrow equivalent) at 6 dpf (fig. S3, A schematic and G). Thus, the excess HSPCs in miR-223Δ/Δ exhibit aberrant proliferation and delamination.

Finally, we examined how miR-223 loss affects blood lineage differentiation of the supernumerary HSPCs. Consistent with the relatively high expression of *miR-223:gfp* in lymphoid/myeloid-primed HSPCs in the AGM (branch 11, Fig. 1D and fig. S2E), lymphoid and myeloid progenitors and differentiated cells were significantly expanded in secondary hematopoietic organs, whereas the erythroid lineage was unchanged (fig. S3H and Fig. 2F). Together, our data suggest that miR-223 limits the production of lymphoid/myeloid-primed nascent HSPCs.

To test if the increased production of hemECs and oligopotent nascent HSPCs is due to loss of miR-223 specifically in ECs, we re-expressed wild-type miR-223 from the endothelial *fli1a* promoter (fig. S3I). miR-223Δ/Δ; *Tg(fli1a:miR-223)* embryos had wild-type numbers of *cmyb+* HSPCs in the AGM and normalization of the lymphoid and myeloid blood cell lineages (Fig. 2, F and G). These data suggest that miR-223 functions in ECs to restrict the EHT of lymphoid/myeloid-primed nascent HSPCs.

Next, we sought to identify the miR-223-dependent mechanism that negatively regulates HSPC production during EHT. miRNAs typically bind miRNA responsive elements (MREs) within 3’ untranslated regions (3’UTRs) to fine tune mRNA levels via decay and/or translational repression (*16*). We identified the endothelial transcripts that were both upregulated upon loss of miR-223 and harbored a predicted miR-223 MRE (*8*). Remarkably, 4 out of the top 8 candidate genes encode enzymes that regulate N-glycosylation (Fig. 3, A and B, fig. S4, A and B, and Data S3).

**Fig 3.**
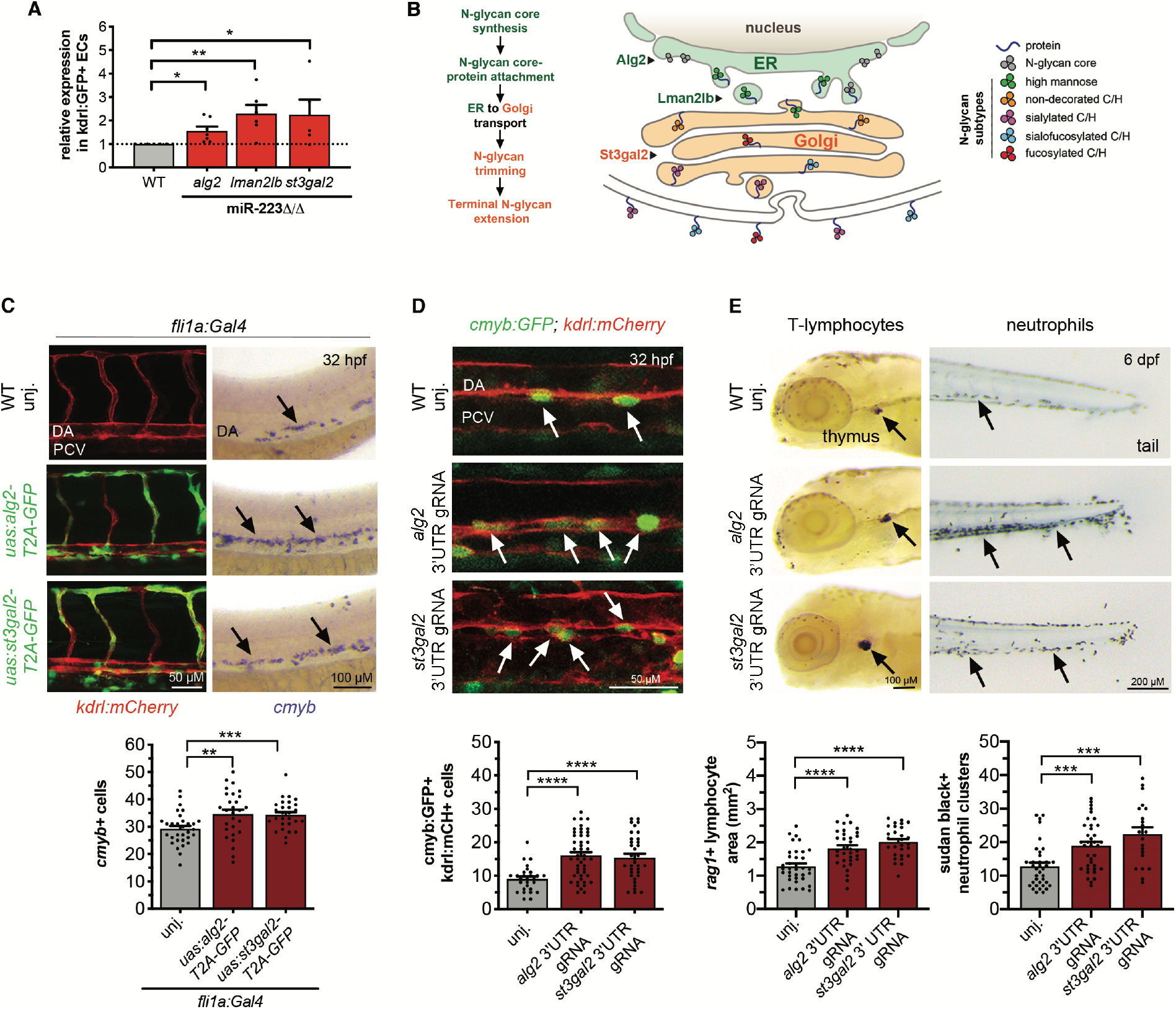
miR-223 limits EHT and lymphoid/myeloid oligopotent HSPC production via EC repression of *alg2* and *st3gal2*. **(A)** Mean ± SEM N-glycogene expression in kdrl:GFP+ ECs at 27 hpf (n = 5-7 biological replicates, paired, two tailed Student’s T-test). **(B)** N-glycan biosynthesis overview. Putative miR-223 target N-glycoenzymes are color-coded based on their function in the ER (green) or Golgi (orange). Arrowheads point to location of N-glycoenzymes functions within the pathway. High mannose and complex/hybrid (C/H) N-glycan subtypes analyzed in Fig. 4 are schematized. **(C)** *top*, Lateral Z-projections (*left)* or brightfield images (*right)* of the zebrafish AGM. GFP marks *alg2* or *st3gal2* overexpression within the kdrl:mCH+ endothelium. *Cmyb* expression labels nascent HSPCs. *bottom*, Mean ± SEM of *cmyb*+ cell number (n = 28 – 31 embryos). **(D)** *top*, Arrows indicate cmyb:GFP+ kdrl:mCh+ nascent HSPCs in the zebrafish AGM. *bottom*, Mean ± SEM of cmyb:GFP+ kdrl:mCH+ cells (n = 28 – 48 embryos). **(E)** *top*, Representative T-lymphocyte marker *rag1* expression and Sudan black neutrophil staining. *bottom*, Mean ± SEM area of *rag1* expression (n = 28 - 35 embryos) and Sudan Black+ clusters (n = 21 - 37 embryos). Significance calculations unless otherwise indicated as in Figure 2. Abbreviations as in Figure 1.

N-glycosylation begins in the endoplasmic reticulum (ER), where a preassembled glycan core is attached *en bloc* onto asparagine (N) residues of nascent polypeptide chains. The glycan is then progressively modified by glycosidases and glycosyltransferases as the protein traffics from the ER to the Golgi (*17*) (Fig. 3B). The rate of glycan flux and/or the expression level of N-glycan biosynthesis enzymes (N-glycoenzymes) produce a diverse N-glycan repertoire, which determines protein activity and important cell behaviors in reprogramming and oncogenesis (*18, 19*). However, N-glycosylation has not yet been implicated in EHT.

We focused on three canonical N-glycoenzymes that were derepressed in miR-223Δ/Δ ECs, namely *alg2*, an ER-resident mannosyltransferase that helps build the N-glycan core by adding mannose moieties; *lman2lb*, a mannose-binding lectin that promotes N-glycoprotein transport from the ER to the Golgi; and *st3gal2*, a Golgi-resident sialyltransferase that terminally modifies N-glycoproteins with sialic acid sugars (Fig. 3, A and B and Data S3).

To determine if miR-223 negatively regulates these N-glycoenzyme transcripts via their 3’UTRs, we constructed sensors consisting of mCherry fused to a 3’UTR fragment of *alg2*, *lman2lb*, or *st3gal2* that contains the putative miR-223 MRE (fig. S4C). The endothelial-specific *fli1a* promoter was used to drive co-expression of both the mCherry sensor and a control eGFP that lacks a 3’UTR (for normalization). Notably, the *alg2* and *st3gal2* sensors showed significantly higher mCherry:eGFP ratios in the AGM of miR-223Δ/Δ versus control embryos (fig. S4C). In contrast, the mCherry:eGFP ratio from the *lman2lb* sensor was similar between wild-type and mutant embryos (fig. S4C). These data suggest that miR-223 represses *alg2* and *st3gal2* transcripts via their 3’UTR in AGM ECs.

We investigated to what extent *alg2* or *st3gal2* repression contributes to miR-223-mediated regulation of EHT and HSPC production. First, we found that like miR-223, *alg2* and *st3gal2* were expressed in ECs, and elevated in nascent HSPCs at 27 hpf (fig. S5A). We used morpholinos to diminish *alg2* or *st3gal2* expression, which significantly reduced nascent HSPCs in the AGM, without otherwise affecting development (fig. S5, B to E). Moreover, partial downregulation of *alg2* or *st3gal2* had no effect on wild-type embryos, but rescued the HSPC expansion in miR-223Δ/Δ embryos (fig. S5F), suggesting that *alg2* or *st3gal2* derepression substantially contributes to the EHT phenotypes observed upon loss of miR-223.

Congruently, *alg2* and *st3gal2* gain-of function models produced phenotypes resembling those in miR-223Δ/Δ embryos. Overexpression of *alg2* and *st3gal2* in wild-type ECs resulted in HSPC expansion in the AGM (Fig. 3C and fig. S6A). Likewise, mutation of the miR-223 MRE within the endogenous *alg2* or *st3gal2* 3’UTR, which is predicted to impair miR-223 binding, led to elevated *alg2* and *st3gal2* levels (fig. S6, B to D), and recapitulated miR-223 Δ/Δ phenotypes: hemECs and HSPCs were expanded in the AGM (Fig. 3D and fig. S6E), as were HSPCs and differentiated lymphoid/myeloid lineage cells within the secondary hematopoietic tissues (Fig. 3E and Fig. S6F). Together, these data suggest that miR-223 post-transcriptionally represses *alg2* or *st3gal2* to regulate EHT.

We next tested if altered N-glycan modifications in ECs might lead to aberrant EHT in miR-223Δ/Δ. Thus, we isolated wild-type or miR-223Δ/Δ kdrl+ ECs during EHT, and used nano-LC/ESI QTOF (*20*), a high resolution mass spectrometry (MS) analytical approach, to profile changes in specific glycan features (Fig. 4A). Glycans were grouped into subtypes, which include high mannose, nondecorated complex/hybrid (C/H), fucosylated C/H, sialylated C/H, and sialofucosylated C/H (Fig. 4B and fig. S7A). Notably, we found that high mannose (~71%) and sialylated C/H (~18%) were the most abundant glycans on proteins from wild-type ECs. In contrast, sialylated C/H (~97%) and sialofucosylated C/H (~3%) were the glycans on proteins from ECs lacking miR-223, concomitant with a remarkable total loss of high mannose type N-glycans (Fig. 4, B and C and Data S4).

**Fig 4.**
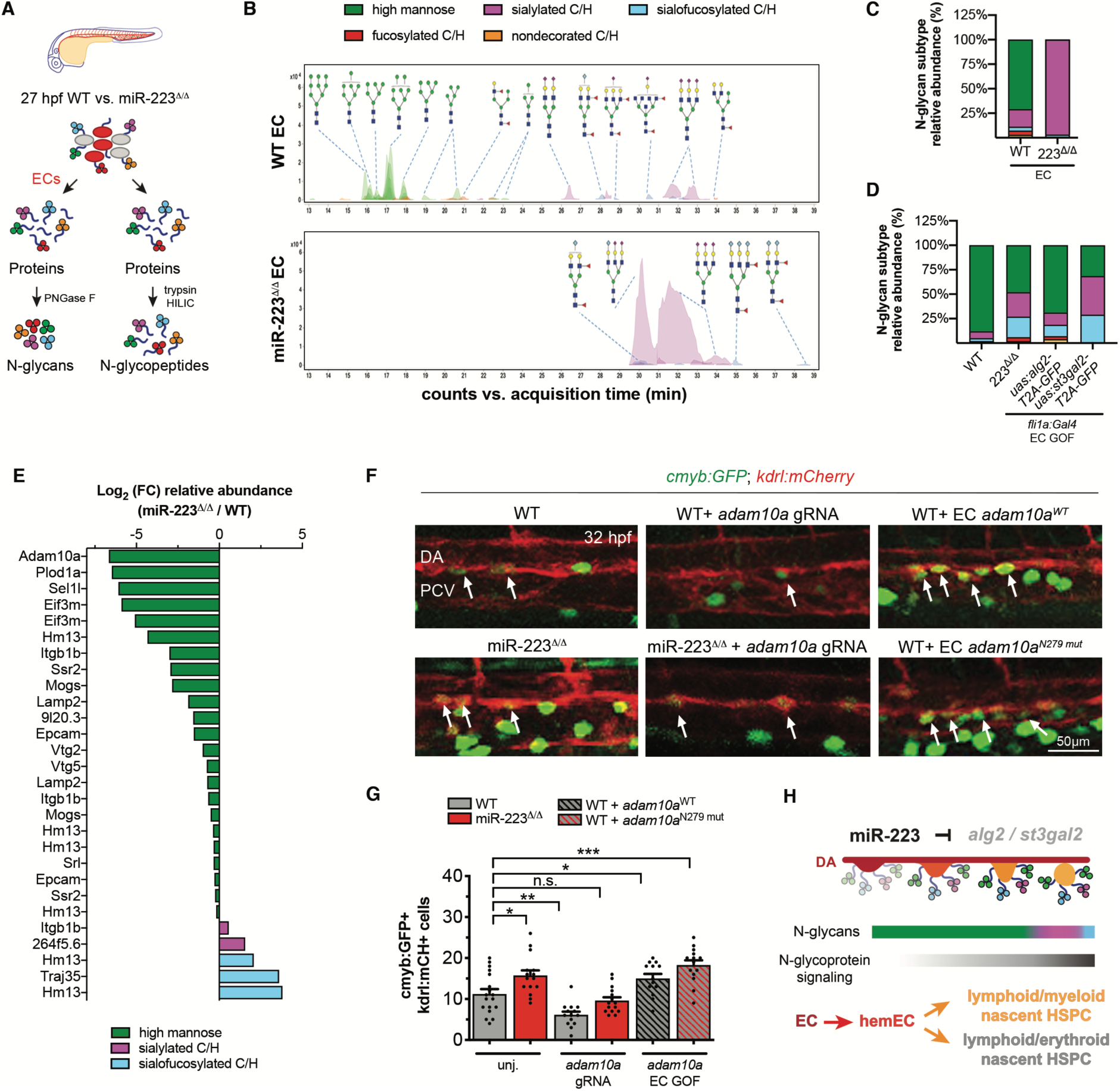
A distinct N-glycan repertoire regulates protein function to limit EHT. **(A)** N-glycan and N-glycopeptide extraction procedure from 27 hpf ECs or whole zebrafish embryos, respectively. Extracted N-glycans or N-glycopeptides were analyzed by MS. **(B)** MS chromatograms for FAC-sorted ECs showing relative abundance of N-glycan subtypes. A subset of the identified N-glycan structures are indicated (see fig. S7A for monosaccharide legend). (**C-D**) Relative abundance of N-glycan subtypes in ECs (**C**) or whole embryos (**D**). N-glycan subtypes are colored as in **B**. **(E)** Log_2_ (fold change) of N-glycan relative abundance for miR-223-regulated N-glycopeptides (n = 3 replicates). **(F)** White arrows point to nascent HSPCs in the zebrafish AGM of embryos injected with a gRNA that diminishes *adam10a* or with EC gain-of function (GOF) *adam10a* constructs. **(G)** Mean ± SEM number of cmyb:GFP+ kdrl:mCH+ HSPCs (n = 13 – 18 embryos). **(H)** Model illustrates how miR-223-dependent regulation of the N-glycome restricts EHT and lymphoid/myeloid oligopotent HSPC production. In ECs, miR-223 represses *alg2* and *st3gal2* N-glycoenzymes, leading to high mannose-(green gradient) versus sialylated-(pink/blue gradients) biased modification of proteins (e.g. Adam10a), which restricts (opacity) their signaling (black gradient). Cell types involved in EHT and N-glycan subtypes are colored according to their defining markers in Fig. 1E or to the legend in **B**, respectively. Significance calculations as in Fig. 2. Abbreviations as in Fig.1

To address whether the altered endothelial N-glycome in miR-223Δ/Δ embryos could result from derepression of the mannosyltransferase *alg2* or the sialyltransferase *st3gal2*, we performed N-glycomic analysis of whole embryos overexpressing *alg2* or *st3gal2* in ECs, which phenocopy miR-223Δ/Δ blood defects (Fig. 3C). Remarkably, we observed a reduction of high mannose and a corresponding increase in sialylated C/H and sialofucosylated C/H modifications, similar to miR-223Δ/Δ embryos (Fig. 4D and Data S4). Thus, our analysis supports that the balance between high mannose versus sialic acid-containing N-glycan subtypes in ECs is regulated by miR-223-mediated repression of *alg2* or *st3gal2* N-glycoenzymes.

Next, we addressed whether compounds that alter the N-glycan repertoire similarly to miR-223 loss could result in an aberrant EHT. Treatment of wild-type embryos with N-Acetylmannosamine (ManNAc), the biological precursor of sialic acid (*21*), increased nascent HSPCs in the AGM, thereby recapitulating the miR-223Δ/Δ phenotype (fig. S7, B and C). Furthermore, N-glycomic analysis of ManNAc-treated embryos confirmed that N-glycoproteins were enriched with sialylated glycan types at the expense of high mannose modfications (fig. S7D). Thus, a distinct N-glycome composition of high mannose to sialylated glycan ratio is necessary to allow normal EHT and HSPC production.

To further investigate how miR-223 regulates EHT via the N-glycome, we identified the N-glycoproteins that were altered in miR-223Δ/Δ compared to wild-type embryos (Fig. 4A). Interestingly, the N-glycoproteins with reduced high mannose and/or increased sialylated C/H and sialofucosylated C/H modifications in miR-223Δ/Δ have known biological functions relevant to EHT and HSPC production and their transcripts were found in trunk ECs at 27 hpf (Fig. 4E, fig. S7E, and Data S4).

Of high interest was the metalloprotease Adam10a, which showed the greatest reduction in high mannose upon miR-223 loss (Fig. 4E) and is critical for the proteolytic release of several cell surface proteins such as Notch, Vegfr2, TNF-α, and other EHT growth factor signaling molecules (*22*–*24*). To determine how miR-223-dependent modification of Adam10a affects EHT, we first used CRISPR/Cas9 to downregulate *adam10a* expression (fig. S7F). We found that the number of nascent HSPCs were significantly decreased in the AGM of *adam10a* gRNA-injected embryos (Fig. 4, F and G), consistent with Adam10-mediated regulation of several EHT factors. Notably, downregulation of *adam10a* in miR-223Δ/Δ embryos decreased nascent HPSC numbers to wild-type levels, suggesting that Adam10a function was enhanced upon miR-223 loss (Fig. 4, F and G). To further support this hypothesis, we overexpressed wild-type *adam10a*, or a mutant version (N279mut), in *fli1a+* ECs (fig. S7, G and H). Mutation of the conserved N279 consensus site prevents N-glycan attachment (*25*). Strikingly, we found that *adam10a* wild-type, and even further the *adam10a* N279mut, exhibited an expansion of nascent HSPCs in the zebrafish AGM (Fig. 4, F and G). These data suggest that decreased high mannose terminal modification of Adam10a promotes its function and contributes to the EHT phenotype of miR-223Δ/Δ embryos.

Our study reveals an unanticipated role for the N-glycan biosynthesis pathway in restricting the production of hemECs, HSPCs, and distinct blood lineages. We discovered an intrinsic mechanism that negatively regulates the transdifferentiation of ECs into oligopotent HSPCs based on a “sugar code” comprised of specific cell surface glycans, high mannose and sialylated complex/hybrid subtypes. This regulation is mediated by miR-223-dependent repression of N-glycan biosynthesis genes, which controls the glycan repertoire to ensure normal HSPC production and differentiation (Fig. 4H and fig. S7I).

Loss of miR-223-mediated regulation of N-glycoenzymes causes a switch in N-glycans and enhances the function of N-glycoproteins. For example, Adam10a EC-autonomous activity is required for EHT, but causes excessive HSPC production when lacking high mannose probably via enhanced regulation of Notch, TNFα and/or other Adam10-dependent signaling pathways.

Furthermore, since EHT is a heterogenous continuum of cells states (*15*), particular hemECs and nascent HSPCs could require miR-223 regulation of N-glycoprotein-dependent signaling to restrict EHT and balance blood production (Fig. 4H and fig. S7I). Congruently, N-glycoproteins throughout the EHT process could influence the differentiation of oligopotent nascent HSPCs. For example, Adam10 gain-of-function during HSPC development effects individual Notch-ligand cleavage products which, in turn, influence cell-fate determination (*22*). Our findings lay the foundation for numerous mechanistic studies of how protein N-glycosylation balances the diverse array of hematopoietic regulators during EHT.

The discovery that EHT is regulated by a specific glycan profile launches new, exciting avenues to explore for the regulation of in vivo and ex-vivo blood stem cell production (*26*). The wide variety of known glycan metabolism modifiers and inhibitors could be used to glyco-engineer hemECs, optimizing the production of glycoforms that facilitate somatic cell reprogramming to HSPCs (*27*). Thus, our findings could inform pharmacological strategies to produce HSPCs for therapeutic interventions.

## Acknowledgments

The authors would like to thank Meredith Cavanaugh for fish husbandry and care, and Sameet Mehta from the Yale Center for Genome Analysis for ScRNA-seq bioinformatic analysis. We would also like to acknowledge Angela Andersen (Life Science Editors), Valentina Greco, Daniel DiMaio, Liana C. Boraas, and Siyuan (Lily) Chen for critical reading of the manuscript. We apologize to those whose original work could not be cited due to space limitations.

## Author Contributions

D.M.K., J.H., and S.N. designed and conducted experiments, analyzed data, and wrote the manuscript. J.G., H.K.M, K.S., and W.A. carried out zebrafish experiments and analyzed data. K.K.H., Y.W. and Z.H. designed and performed mouse phenotypic experiments and analyses. C.L., Y.S. and Y. X. conducted glycomic and glycoproteomic analyses. L.K.M., D.W.H., W.C.S., and EJ. P. contributed N-glycome-related expertise and preliminary data. All authors edited the paper.

## Data and materials availability

Previously published bulk cell RNA-seq experiments are submitted to the Gene Expression Omnibus (http://www.ncbi.nlm.nih.gov/geo/) under accession number GEO: GSE81341 and raw Quant-seq reads are publicly accessible in the Sequence Read Archive under SRP099466. Single cell RNA-seq experiments generated for this study are submitted under accession number GEO:GSE135246.

## Supplementary Materials

Materials and Methods

Figures S1-S7

Movies S1-S3

Data S1-S4

References

